# *In silico* epitope prediction and 3D model analysis of Peste des petits ruminants virus nucleoprotein (PPRV N)

**DOI:** 10.1101/095505

**Authors:** Bakang Baloi

## Abstract

Peste des petits ruminants virus (PPRV) is an acute, highly contagious viral disease of small ruminants. It is endemic to sub-Saharan Africa, Asia and the Arabian Peninsula. The disease is a major constraint to food security, causing significant economic losses to subsistence farmers in affected areas.

The nucleoprotein of morbilliviruses is highly immunogenic and produced in large quantities in virus infected cells. This makes it a suitable target for the host’s immune response. In this study, B-cell and T-cell epitopes of PPRV Nig/75/1 nucleoprotein were predicted using a suite of *in silico* tools. Forty-six T-cell epitopes were predicted, of which 38 were MHC-I binding while eight were MHC-II binding. Of the 19 B-cell epitopes predicted, 15 were linear epitopes while four were discontinuous epitopes. Homology modelling of PPRV-N was done to elucidate the 3D structure of the protein and conformational epitopes. Conservation analysis of the discontinuous epitopes gave an indication into the similarity of the selected epitopes with other isolates of PPRV.

Predicted epitopes may form an important starting point for serological screening and diagnostic tools against PPRV. Experimental validation of the predicted epitopes will assist in selection of promising candidates for consideration as antigen-based diagnostic tools. Such diagnostic tools would play a role in the global fight and possible eradication of PPR.

## Introduction

Peste des petits ruminants virus (PPRV) highly contagious transboundary viral disease of goats and sheep widely distributed in sub-Saharan Africa and the Arabian Peninsula (Shaila *et al.,* 1996). The disease causes significant economic losses, especially to subsistence farmers who do not have easy access to vaccination. It is considered one of the main constraints to small ruminants’ production and is listed as a reportable disease by the Office International des Epizooties (OIE) (Berhe *et al.,* 2003). Morbidity and mortality rates vary but can be as high as 100% and 90%, respectively in immunologically naive populations (Pope *et al.,* 2013). The disease is caused by Peste des petit ruminant virus (PPRV); a single-stranded, negative sense RNA virus of the genus Morbillivirus, in the family Paramyxoviridae (Diallo *et al.,* 1994).

The nucleoprotein of morbilliviruses is the most abundant structural protein and an important regulator of replication and transcription (Ismail *et al.,* 1995). It is highly immunogenic in spite of its internal location (Yu *et al.,* 2015) and expressed to a very high level in morbillivirus-infected cells (Choi *et al.,* 2004). It consists of 525 amino acids with an estimated molecular weight of 58 kDa. (Diallo *et al.,* 1994). After infection anti-N antibodies are produced indicating that there is a direct release of morbillivirus nucleoprotein into the extracellular compartment, where it binds to B-cell receptors (Bodjo *et al.,* 2007). The nucleoprotein protein is therefore a good antigen candidate for the development of differential tests for differentiating infected animals from ones vaccinated (DIVA) with fusion (F) or haemagglutinin (H) based-recombinant marker vaccines (Choi *et al.,* 2004; Choi *et al.,* 2005). As a result the N protein is suitable for serologic screening of naturally infected from vaccinated animals (Couacy-Hymann *et al.,* 2002) and as a diagnostic antigen towards which antibodies are directed during infection (Dechamma *et al.,* 2006). N protein-specific T cells were previously found to comprise the bulk of the virus specific memory cells in the paramyxovirus family (Mitra-Kaushik *et al.,* 2001). To underscore the importance of N protein in the immune response against morbilliviruses, most epitope mapping has been on the nucleoprotein protein (Mitra-Kaushik *et al.,* 2001; Choi *et al.,* 2004; Choi *et al.,* 2005; Dechamma *et al.,* 2006; Bodjo *et al.,* 2007; Yadav *et al.,* 2009; Yu *et al.,* 2015). Though suspected to be unimportant for humoral protection against morbillivirus infection (Couacy-Hymann *et al.,* 2002), the N protein is crucial in development of PPRV serological tests for diagnosis and disease surveillance (Yadav *et al.,* 2009). Descriptions of PPRV N epitopes have been made for B-cell epitopes (Choi *et al.,* 2005; Bodjo *et al.,* 2007) and for both T-helper and B-cell epitopes (Dechamma et al., 2006).

In this study, the 3D model of PPRV nucleoprotein (also known as PPRV gP1) was determined and an integrated *in silico* approach used to predict both B-cell and T-cell epitopes. The predicted epitopes may form an important starting point for serological screening and diagnostic tools against PPRV.

## Materials and methods

### Retrieval of protein sequences and conservation analysis

All available full amino acid sequences of the PPRV nucleoprotein (PPRV N) were retrieved from the National Center for Biotechnology Information (NCBI) (http://www.ncbi.nlm.nih.gov/nucleotide) database in FASTA format. Conserved regions of PPRV N were determined with the web version of Clustal Omega (McWilliam *et al.,* 2013) multi sequence alignment (MSA) tool from the European Bioinformatics Institute (EMBL-EBI) (http://www.ebi.ac.uk/Tools/msa/clustalo/) in default parameters. Redundant sequences were excluded from the alignment. Even though an NCBI reference sequence for PPRV nucleoprotein exists: accession number (YP_133821.1) (Bailey *et al.,* 2005), the sequence for PPRV strain Nigeria/75/1, GenBank accession number: CAA52454.1 (Diallo *et al.,* 1994) was the one used throughout this study based on locally available technical support with the particular strain.

### T cell epitope prediction MHC class I prediction

MHCI binding predictions were made with artificial neural networks (ANNs) based methods such as the allele specific NetMHC (Lundegaard *et al.,* 2008) and pan-specific NetMHCpan (Hoof *et al.,* 2009). NetMHC results were obtained from the NetMHC 4.0 Server (Nielsen and Andreatta, 2016), accessed from (http://www.cbs.dtu.dk/services/NetMHC/) while NetMHCpan results were obtained from NetMHCpan version 3.0 server (http://www.cbs.dtu.dk/services/NetMHCpan/). Predictions were made based on the selection of between 9mer and 14mer mouse alleles: H-2-Db,H-2-Dd,H-2-Kb,H-2-Kd, H-2-Ld, H-2-Qa1 and H-2-Qa2 with threshold for strong binders set at 0.5 and for weak binders at 2. Cross binding epitopes were selected as putative epitopes.

Furthermore, proteosomal processing predictors based upon neural network architecture such as NetChop (Kesmir *et al.,* 2002; Saxova *et al.,* 2003; Nielsen *et al.,* 2005), NetCTL (Larsen *et al.,* 2005; Larsen *et al.,* 2007) and NetCTLpan (Stranzl *et al.,* 2010) were utilised. Epitope candidates based on peptide processing within the cell were determined from the IEDB website (http://tools.iedb.org/main/tcell/). Parameters selected for NetChop analysis were C terminal (3.0) with a 0.5 threshold. For NetCTL the weight on C terminal cleavage was set at 0.15, weight on TAP transport efficiency 0.05, selected supertype B7 and threshold varied between 0.2 and 0.75. For NetCTLpan (NetCTLpan 1.1 Server; http://www.cbs.dtu.dk/services/NetCTLpan/) epitope candidates were predicted for different mouse alleles based on the following settings: weight on C terminal cleavage (0.225), weight on TAP transport efficiency (0.025), threshold for epitope identification (1.0), threshold for showing predictions (-99.9) High scoring epitopes recognised by multiple alleles were selected.

### MCH class II prediction

MHCII binding predictions for mouse H-2-I locus (H2-IAd and H2-IEd alleles) were made using the IEDB analysis resource Consensus tool (Wang *et al.,* 2008; Wang *et al.,* 2010). This tool combines any three of the four methods; NN-align (Nielsen and Lund, 2009), SMM-align (Nielsen *et al.,* 2007), Sturniolo (Sturniolo *et al.,* 1999) and NetMHCIIpan (Nielsen *et al.,* 2008) to predict MHC Class II epitopes. Strong binding peptides and those binding to multiple MHC class II molecules were selected.

### B-cell epitope predictions Prediction of linear B-cell epitopes from antigen sequence properties

A variety of tools were used to predict linear epitopes along the 525 amino acid PPRV N protein. IEDB B-cell epitope prediction tools used included BepiPred (Larsen *et al.,* 2006), Chou and Fasman beta-turn prediction (Chou and Fasman, 1979), Emini accessibility prediction (Emini *et al.,* 1985), Karplus and Schulz flexibility prediction (Karplus and Schulz, 1985), Kolaskar and Tongaonkar antigenicity prediction method (Kolaskar and Tongaonkar, 1990) and Parker hydrophilicity plot (Parket *et al.,* 1986). B-cell epitope prediction methods; BCPred (EL-Manzalawy *et al.,* 2008) and implementation of the AAP method (Chen *et al.,* 2007) from the BCPREDS server (http://ailab.ist.psu.edu/bcpred/) were also used. Results acquired from the BCPREDS server were compared with BepiPred obtained results. Epitopes with high scores, and those common between different prediction methods were selected as likely to be antigenic.

### Structural epitope prediction

Structural predictions of B-cell epitopes were done with the ElliPro antibody epitope prediction tool (http://tools.iedb.org/ellipro/). ElliPro, is a web-tool that allows the prediction and visualization of antibody epitopes in a given protein sequence or structure (Ponomarenko *et al.,* 2008). Structural epitopes were generated from the PDB file of PPRV N 3D protein model generated by Phyre2 (Kelley *et al.,* 2015). Epitope structure predictions were performed based on default parameters (minimum score value 0.5 and maximum distance of 6Å). High scoring three dimensional epitope structures were viewed with the Jmol applet (http://www.jmol.org/).

### 3D Modelling of PPRV N

Searching the Protein Databank Archive (RCSB PDB) (http://www.rcsb.org/pdb/home/home.do) revealed that there were no records of 3D structures for PPRV N. The three dimensional protein structure of PPRV N was predicted with the Protein Homology/analogY Recognition Engine V 2.0 (Phyre2 V2.0) (Kelley *et al.,* 2015) server (http://www.sbg.bio.ic.ac.uk/phyre2/html/page.cgi?id=index) in ‘Intensive’ modelling mode. Phyre2 uses advanced remote homology detection methods to build 3D models, predict ligand binding sites and analyse the effect of amino acid variants in protein sequences (Kelley *et al.,* 2015).

### Ligand prediction

The ligand binding site on the modelled 3D structure of PPRV N was predicted with 3DLigandSite (http://www.sbg.bio.ic.ac.uk/3dligandsite). 3DLigandSite is a high performing web server for automatic prediction of ligand-binding sites for protein that have not been solved (Wass *et al.,* 2010).

### Protein structure quality and validation

The resulting 3D model was analysed and viewed with Swiss-PdbViewer Deep View Version 4.1 (http://spdbv.vital-it.ch/) and the Jmol viewer (http://www.jmol.org/). Different tools were also employed for stereochemical analyses and model quality evaluation. Structural evaluation and validation were performed using the Structure Analysis and Verification Server (http://services.mbi.ucla.edu/SAVES/) to obtain ERRAT (Colovos and Yeats, 1993), VERIFY 3D (Bowie *et al.,* 1991) and PROVE (Pontius *et al.,* 1996) scores. In addition Ramachandran plots of the model were generated using the RAMPAGE web tool (Lovell *et al.,* 2002) available at (http://mordred.bioc.cam.ac.uk/˜rapper/rampage.php).

## Results and discussion MHC prediction

T-cell recognition of peptide/major histocompatibility complex (MHC) is a prerequisite for cellular immunity. Different MHC molecules bind peptide fragments from pathogens and in association with T-cell receptor (TCR) molecules present them to appropriate T-cells (Zepp,2016). Peptide fragments that bind specific MHC are therefore targets for vaccine and immunotherapy (Tong *et al.,* 2007). As such their accurate prediction is important in the design of diagnostics and vaccines. MHC I binds and presents epitopes which are derived from proteolytically degraded intracellular proteins about 8–11 residues long while MHC II epitopes are derived from extracellular sources and are much longer on average (up to 25 residues long) (Michalik *et al.,* 2016). Differences in the structures of MHC molecules mean that peptides bind differently to the binding pockets of each MHC class (Wang *et al.,* 2008). Different prediction tools may therefore be required to exploit differences in structure and binding affinities of the two MHC classes. MHC class I epitopes (Tables 1 and 2; Figure 1) and MHC class II epitopes (Table 3) were predicted using NetMHC, NetCTLpan, NetCHOP and NetCTL. Selected epitopes were those binding multiple alleles (Tables 1 and 2) and those taken to be strongly binding (Table 3) based on binding threshold. Most epitopes described as either binding different types of alleles, or strong binders (less than 2% binding threshold) corresponded with epitopes and minimal motifs described by Yu *et al.,* (2015) through fine mapping and conservation analysis. The distribution of epitopes within the nucleoprotein is graphically displayed in Figure 1. The observed overlap between predicted and experimentally determined epitopes implies a high likelihood that described epitopes are antigenic.

**Figure 1.**
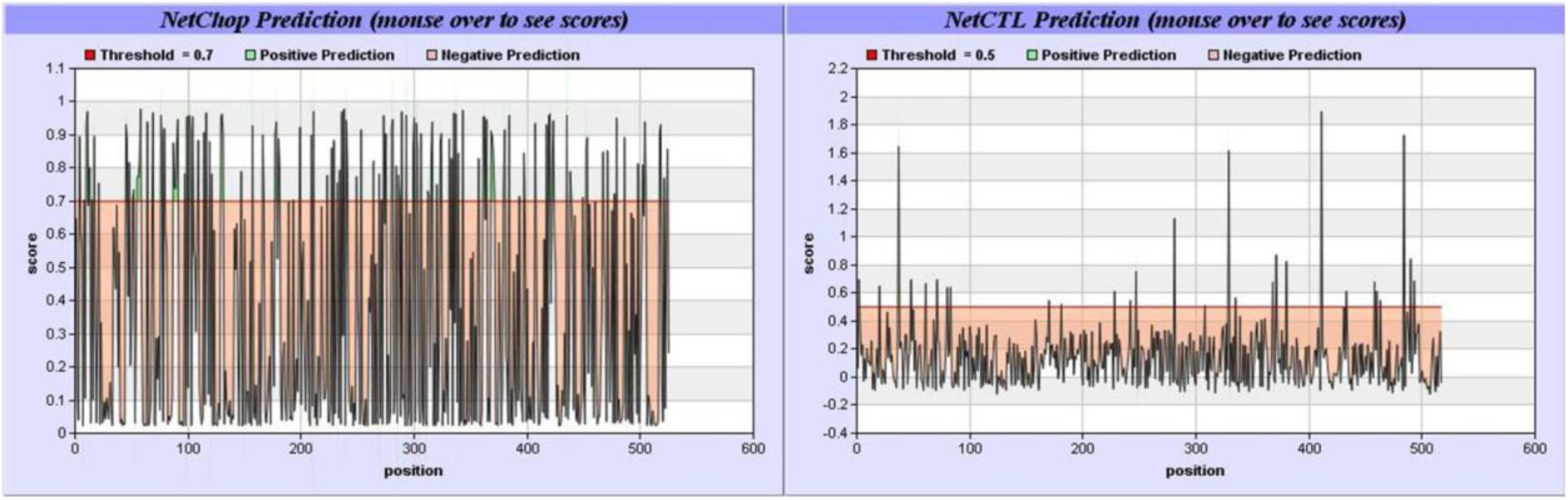
Graphical representation of NetCHOP and NetCTL predicted PPRV N epitopes.

**Table 1.**
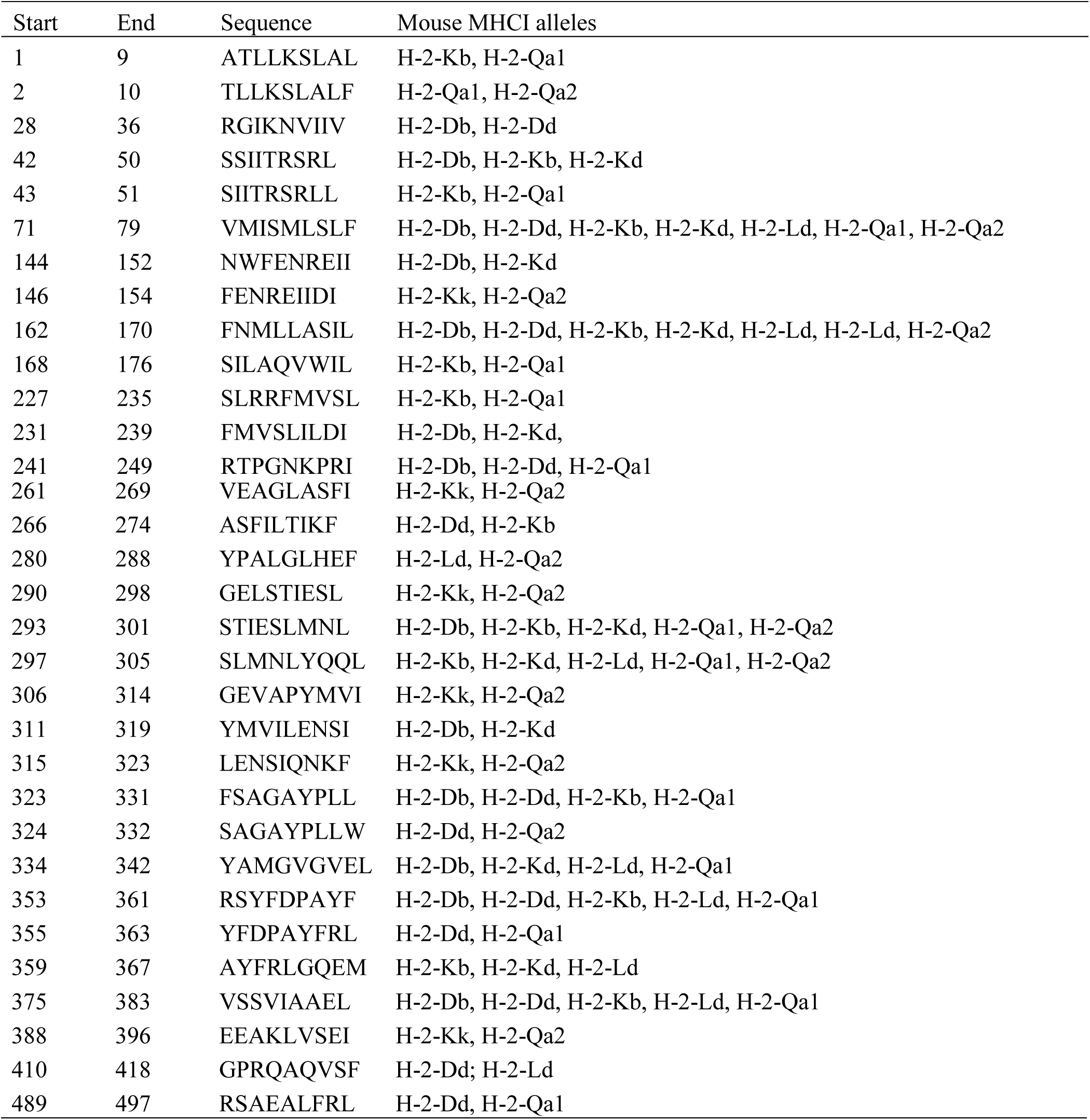
NetMHC and predicted cross-binding epitopes of PPRV nucleoprotein.

**Table 2.** NetCTLpan predicted promiscuous epitopes.

| Position | Peptide sequence | Allele |
| --- | --- | --- |
| 0 | MATLLKSLALF | H-2-Db H-2-Dd H-2-Kb H-2-Kd H-2-Kd H-2-Kk H-2-Ld |
| 6 | SLALFKRNKDK | H-2-Kd H-2-Kk |
| 47 | RSRLLDRLVRL | H-2-Db H-2-Dd H-2-Kd H-2-Kk H-2-Ld |
| 53 | RLVRLAGDPDI | H-2-Db H-2-Dd H-2-Kk H-2-Ld |
| 80 | VESPGQLIQRI | H-2-Db H-2-Dd H-2-Ld |
| 105 | QSTRSQSGLTF | H-2-Db H-2-Dd H-2-Ld |

**Table 3.** NetMHCIIpan predicted strongly binding MCH II epitopes of PPRV nucleoprotein.

| Position | Peptide sequence | Interacting allele | % Rank (Threshold for strong binding peptides- 2%) |
| --- | --- | --- | --- |
| 126 | ADMYFSTEGPSSGSK | H-2-Iab | 1.6 |
| 434 | REEVKAAIPNGSEGR | H-2-Iab | 1.9 |
| 127 | DMYFSTEGPSSGSKK | H-2-Iab | 2 |
| 96 | VSIRLVEVVQSTRSQ | H-2-Iad | 0.2 |
| 492 | EALFRLQAMAKILED | H-2-Iad | 0.8 |
| 173 | VWILLAKAVTAPDTA | H-2-Iad | 1.3 |
| 387 | AEEAKLVSEIASQTG | H-2-Iad | 1.5 |
| 226 | LSLRRFMVSLILDIK | H-2-Iad | 1.7 |

**Table 4.**
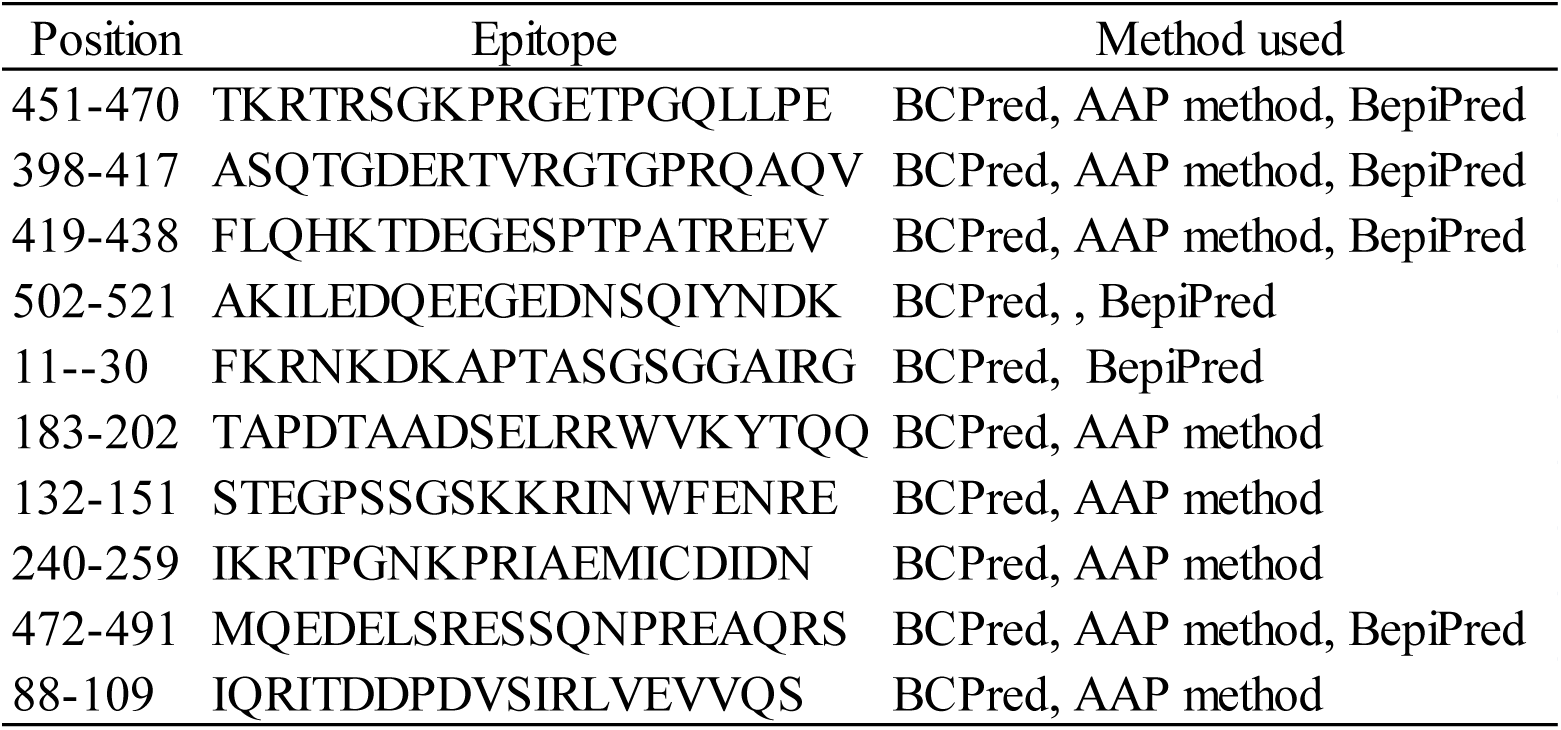
B-cell epitopes predicted by BepiPred related methods.

## Prediction of linear B-cell epitopes from PPRV N sequence properties

The humoral immune response recognises antigenic determinants in pathogenic proteins to activate and generate memory B-cells. These antigenic regions are called B-cell epitopes and can be used to develop diagnostic tools and vaccines against pathogens (Dhanda *et al.,* 2016). Consequently, reliable prediction of antibody, or B-cell epitopes is important in the design of vaccines and immunodiagnostics (Ponomarenko *et al.,* 2008). Unlike T-cell epitopes, B-cell epitopes are not presented in the context of MHC molecules (Michalik *et al.,* 2016). Additionally, they often exist as discontinuous epitopes and are known to be difficult to predict (Kringelum *et al.,* 2012). Tools for identifying B-cell epitopes rely on primary sequence information and functional characteristics of the epitope (Ponomarenko and van Regenemortel, 2009; Sun *et al.,* 2015). This is in contrast with sequence-based methods which rely on descriptors such as ability to form linear secondary structures, hydrophilicity (Hopp and Woods, 1981), hydrophobicity (Kyte and Doolittle, 1982), surface accessibility (Emini *et al.,* 1985). Methods which depend on amino acid composition are accurate for continuous epitope prediction (Ponomarenko and van Regenmortel, 2009).

The methods employed in this study take advantage of various amino acid properties to determine areas of the protein likely to be antigenic. BepiPred predictions for linear B-cell epitopes (Table 3, Figure 2A) rely on a combination of hidden Markov models and propensity scales. The BepiPred method has been found to perform better than both random predictions and propensity scale methods (Larsen *et al.,* 2006).

**Figure 2.**
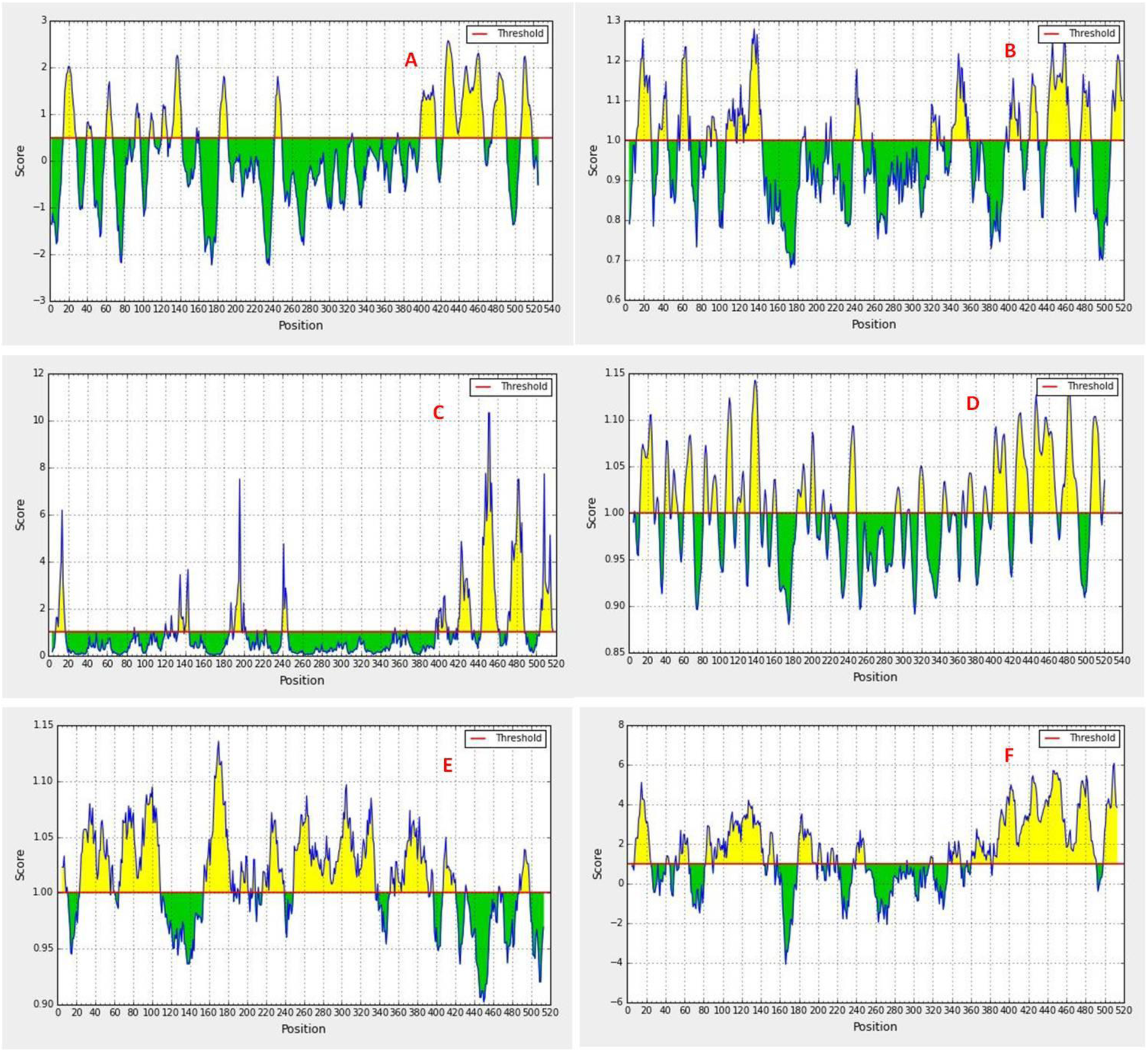
PPRV N epitopes predicted based on sequence properties. A) BepiPred, B) Chou and Fasman beta-turn prediction, C) Emini accessibility prediction D) Karplus and Schulz flexibility prediction, E) Kolaskar and Tongaonkar antigenicity prediction method, F) Parker hydrophilicity plot.

A comparison of sequence based methods for epitope prediction (Figure 2A-2E) indicates that the different methods predict similar areas of the PPRV nucleoprotein as likely to be antigenic. This overlap in epitope prediction increases the chances that identified epitopes will provide the required immune response for vaccine design, or binding specificity in diagnostics. Chou and Fasman beta-prediction method (Figure 2B) uses automated prediction of the chain reversal regions of globular proteins using bend frequencies and β-turn conformational parameters (Chou and Fasman, 1979).

Emini accessibility prediction (Figure 2C) compares sequences by predicted surface features based upon indices of surface probability (Emini *et al.,* 1985). Karplus and Schulz flexibility plot (Figure 2D) takes chain flexibility as indicative of an antigenic determinant and as a basis for selecting cross reacting peptides (Karplus and Schulz, 1985).

Epitopes predicted by the Kolaskar and Tongaonkar antigenicity prediction method (Figure 2E) are based on physicochemical properties of amino acid residues and their frequencies of occurrence in experimentally known segmental epitopes (Kolaskar and Tongaonkar, 1990). Applying this method to a large number of proteins has been shown to perform better than most of the known methods; predicting antigenic determinants by up to 75% accuracy (Kolaskar and Tongaonkar, 1990).

Epitopes predicted by the Parker hydrophilicity method (Figure 2F) were also similar to those predicted by other methods used in the study, or previously described in the literature (Yu *et al.,* 2015). Parker hydrophilicity method is based on peptide retention times during high-performance liquid chromatography (HPLC) in which HPLC parameters showed high correlation with antigenicity (Parket *et al.,* 1986).

B-cell epitopes predicted in this study compare favourably with morbillivirus epitopes mapped by Bodjo *et al.,* (2007) who described Rinderpest virus epitopes that mapped to an area between amino acids 120 and 149 and in the carboxy terminal region between aa 421-525 (Bodjo *et al.,* 2007). Similarly, Choi *et al.,* (2004) described morbillivirus epitopes in the aa regions 1-149, 440-452 and 479-486 which are reflected in figures 3A, 3B, 3C, 3D and 3F of this study (Choi *et al.,* 2004). Based on the conserved nature of the morbillivirus nucleoprotein, and the immunological similarity of both rinderpest and PPRV it is possible that the epitopic regions may occur on the same protein regions. PPRV specific N protein epitopes (Choi *et al.,* 2005; Dechamma *et al.,* 2006) were also described within the amino acid regions 1-262 and 448-521. The advantage of *in silico* methods over experimental methods is that *in silico* methods are quicker and span the entire genome making it easier to predict antigenic areas not recognisable as epitopic experimentally.

### Structural epitope prediction

Sequence-based epitope prediction methods are inadequate in identifying discontinuous B-cell epitopes (Michalik *et al.,* 2016). Discontinuous epitopes are best described by structural methods which incorporate the peptides’ three dimensional structure (Kringelum *et al.,* 2012) and functional properties of the epitope (Sun *et al.,* 2015). In this study, the structure based tool ElliPro was used to predict linear and discontinuous epitopes of PPRV N. ElliPro (from **Elli**psoid and **Pro**trusion), utilises three algorithms to approximate the protein shape as an ellipsoid; calculate the residue protrusion index (PI); and cluster neighbouring residues based on their PI values (Ponomarenko *et al.,* 2008). Five PPRV N epitopes with protrusion indices (PI) above 0.7 (Table 5) were selected and Jmol visualisation of the epitope residues shown in Figure 3 (A-E). A similar approach was used for discontinuous epitopes, where epitopes with PI scores above 0.6 were selected. Results are shown in Table 6, with visualisation of the structures in Figure 4A-D. In both instances, the epitopes are similar to those described by other methods showing the reliability of combined methods in predicting possible antigenic sites. Conservancy results for the epitopes (Tables 7 and 8) show the sequence identity of the epitopes to various isolates and accessions of PPR. Conservancy ranged from about 24% to almost 90% of the 49 PPRV strains for linear epitopes (Table 7). For discontinuous epitopes (Table 8) conservancy ranged from 16% to 98%. Conserved sequences within different strains are possibly in parts of the protein which are indispensable for protein function and not prone to mutations (Diallo *et al.,* 1994). Epitopes from the pathogen’s conserved protein sequences are likely to be effective for various isolates of the pathogen (Sette *et al.,* 2001). Epitopic residues with high sequence identity and high PI score are likely to be antigenic for different PPRV strains, and therefore suitable for further analysis.

**Figure 3.**
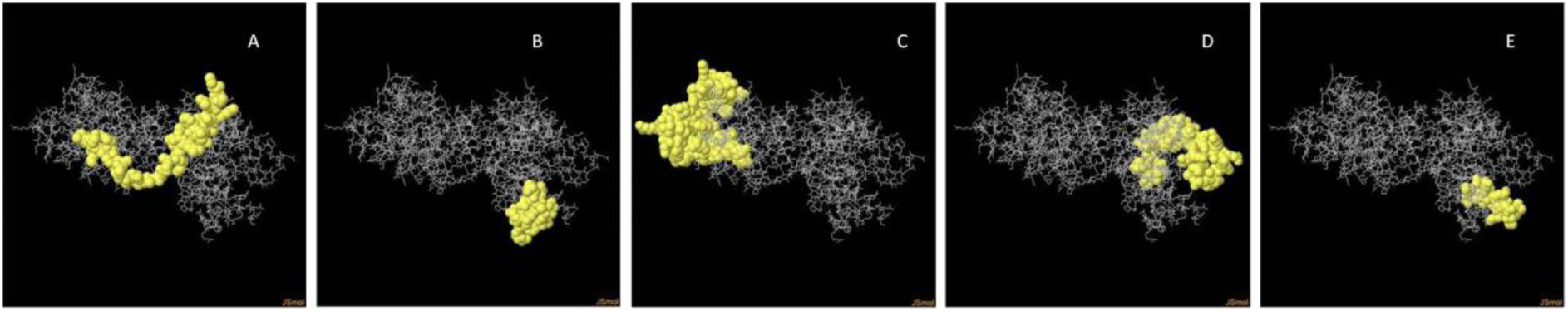
3D depictions of ElliPro predicted linear PPRV N epitopes.

**Figure 4.**
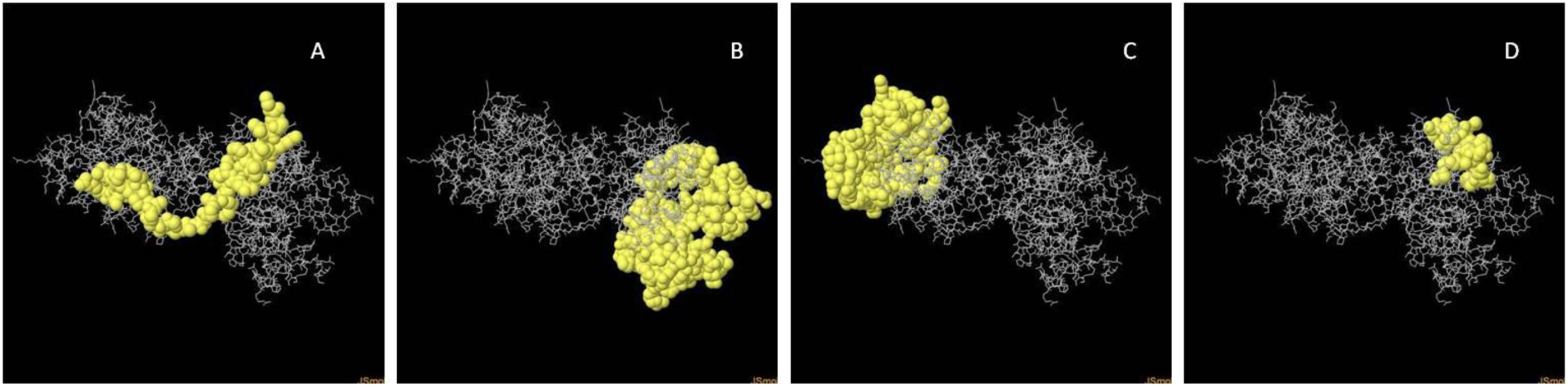
3D depictions of ElliPro predicted discontinuous PPRV N epitopes 3D Modelling of PPRV N.

**Table 5.**
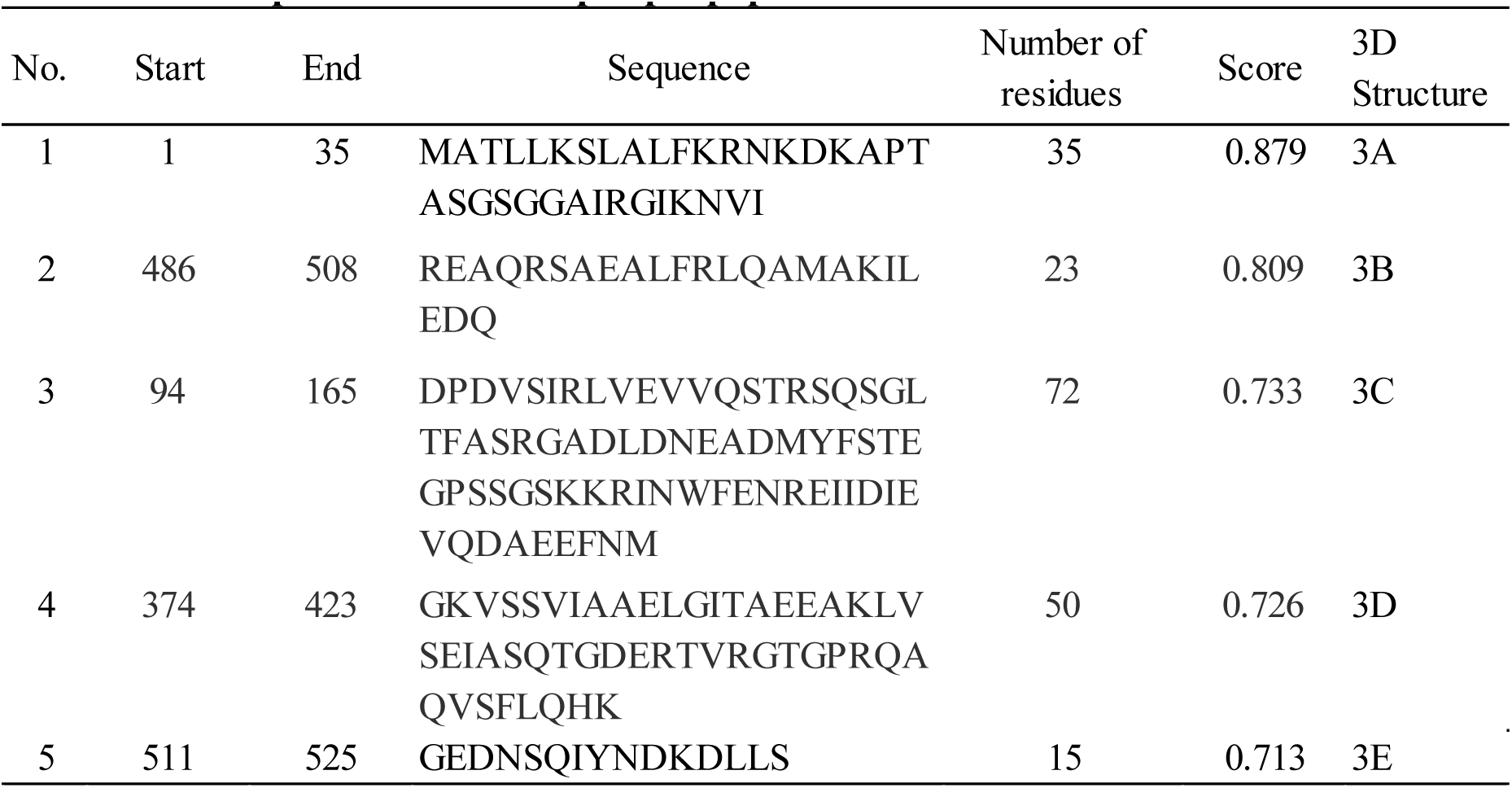
ElliPro predicted linear epitopic peptides of PPRV N.

**Table 6.** ElliPro predicted discontinuous epitopic peptides of PPRV N.

| No. | Residues | Number of | Score | 3D Structure |
| --- | --- | --- | --- | --- |
| 1 | _M1, _A2, _T3, _L4, _L5, _K6, _S7, _L8, _A9, _L10, _F11, _K12, _R13, _N14, _K15, _D16, _K17, _A18, _P19, _T20, _A21, _S22, _G23, _S24, _G25, _G26, _A27, _I28, _R29, _G30, _I31, _K32, _N33, _V34, _I35, _D94, _P95, _D96, _V97, _S98, _I99, _R100 | 42 | 0.824 | 4A |
| 2 | _N345, _E367, _R370, _R371, _G374, _K375, _S377, _S378, _V379, _I380, _A381, _A382, _E383, _L384, _G385, _I386, _T387, _A388, _E389, _E390, _A391, _K392, _L393, _V394, _S395, _E396, _I397, _A398, _S399, _Q400, _T401, _G402, _D403, _E404, _T406, _V407, _R408, _G409, _T410, _G411, _P412, _R413, _Q414, _A415, _Q416, _V417, _S418, _F419, _L420, _Q421, _H422, _K423, _P430, _T431, _P432, _A433, _T434, _R435, _E436, _E437, _V438, _E462, _T463, _P464, _G465, _Q466, _L467, _L468, _P469, _E470, _I471, _M472, _R479, _E480, _S481, _S482, _N484, _E487, _A488, _Q489, _R490, _S491, _A492, _E493, _A494, _L495, _F496, _R497, _L498, _Q499, _A500, _M501, _A502, _K503, _I504, _L505, _E506, _D507, _Q508, _E509, _E510, _G511, _E512, _D513, _N514, _S515, _Q516, _I517, _Y518, _N519, _D520, _K521, _D522, _L523, _L524, _S525 | 116 | 0.702 | 4B |
| 3 | _P40, _G41, _D42, _S43, _S44, _I45, _I46, _T47, _R48, _S49, _R50, _D53, _R54, _R57, _G60, _D61, _P62, _D63, _I64, _L79, _L101, _V102, _E103, _V104, _V105, _Q106, _S107, _T108, _S110, _Q111, _S112, _G113, _L114, _T115, _F116, _A117, _S118, _R119, _G120, _A121, _D122, _L123, _D124, _N125, _E126, _A127, _D128, _M129, _F131, _S132, _T133, _E134, _G135, _P136, _S137, _S138, _G139, _S140, _K141, _K142, _R143, _I144, _N145, _W146, _F147, _E148, _N149, _E151, _I152, _I153, _D154, _I155, _E156, _V157, _Q158, _D159, _A160, _E161, _E162, _F163, _M165, _R204, _V205, _I206, _G207, _E208, _F209, _R210, _L211, _D212, _K213 | 91 | 0.702 | 4C |
| 4 | _Y281, _P282, _A283, _G285, _L286, _H287, _E288, _F289, _A290, _G291, _S294, _L331, _F357, _D358, _P359, _A360, _Y361, _R363 | 18 | 0.638 | 4D |

**Table 7.**
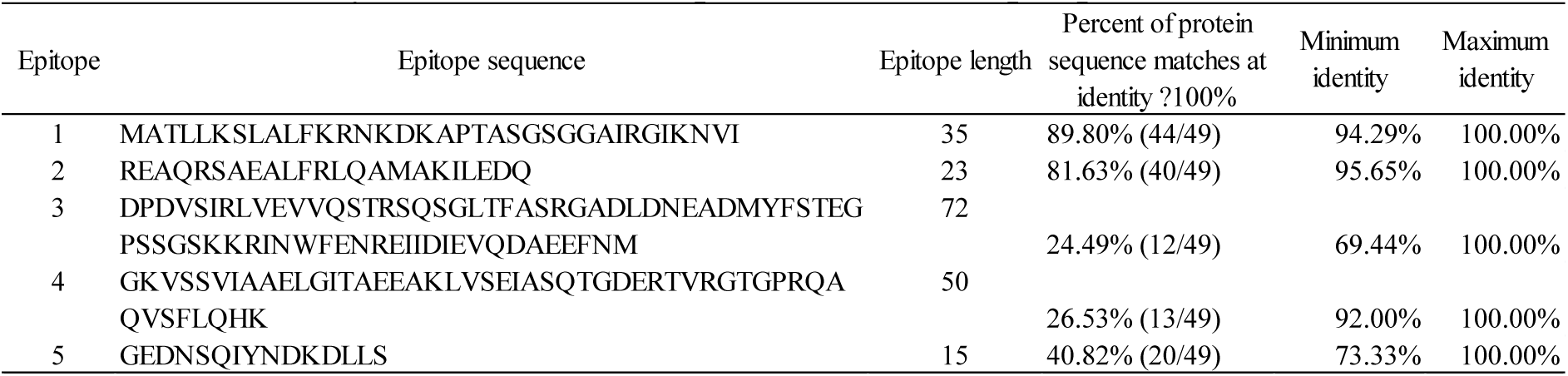
Conservancy result for ElliPro predicted linear epitopes.

**Table 8.** Conservancy result for ElliPro predicted discontinuous epitopes.

| Epitope | Epitope sequence | Epitope length | Percent of protein sequence matches at identity ?100% | Minimum identity | Maximum identity |
| --- | --- | --- | --- | --- | --- |
| 1 | M1, A2, T3, L4, L5, K6, S7, L8, A9, L10, F11, K12, R13, N14, K15, D16, K17, A18, P19, T20, A21, S22, G23, S24, G25, G26, A27, I28, R29, G30, I31, K32, N33, V34, I35, D94, P95, D96, V97, S98, I99, R100 | 42 | 89.80% (44/49) | 95.24% | 100.00% |
| 2 | N345, E367, R370, R371, G374, K375, S377, S378, V379, I380, A381, A382, E383, L384, G385, I386, T387, A388, E389, E390, A391, K392, L393, V394, S395, E396, I397, A398, S399, Q400, T401, G402, D403, E404, T406, V407, R408, G409, T410, G411, P412, R413, Q414, A415, Q416, V417, S418, F419, L420, Q421, H422, K423, P430, T431, P432, A433, T434, R435, E436, E437, V438, E462, T463, P464, G465, Q466, L467, L468, P469, E470, I471, M472, R479, E480, S481, S482, N484, E487, A488, Q489, R490, S491, A492, E493, A494, L495, F496, R497, L498, Q499, A500, M501, A502, K503, I504, L505, E506, D507, Q508, E509, E510, G511, E512, D513, N514, S515, Q516, I517, Y518, N519, D520, K521, D522, L523, L524, S525 | 116 | 18.37% (9/49) | 87.07% | 100.00% |
| 3 | P40, G41, D42, S43, S44, I45, I46, T47, R48, S49, R50, D53, R54, R57, G60, D61, P62, D63, I64, L79, L101, V102, E103, V104, V105, Q106, S107, T108, S110, Q111, S112, G113, L114, T115, F116, A117, S118, R119, G120, A121, D122, L123, D124, N125, E126, A127, D128, M129, F131, S132, T133, E134, G135, P136, S137, S138, G139, S140, K141, K142, R143, I144, N145, W146, F147, E148, N149, E151, I152, I153, D154, I155, E156, V157, Q158, D159, A160, E161, E162, F163, M165, R204, V205, I206, G207, E208, F209, R210, L211, D212, K213 | 91 | 16.33% (8/49) | 57.75% | 100.00% |
| 4 | Y281, P282, A283, G285, L286, H287, E288, F289, A290, G291, S294, L331, F357, D358, P359, A360, Y361, R363 | 18 | 97.96% (48/49) | 94.44% | 100.00% |

The 3D model of PPRV N was predicted with Phyre2 in the ‘Intensive’ mode. Phyre2 is a homology modelling tool for prediction and analysis of protein structure, function and mutations. Phyre2 uses Hidden Markov Models (HMM) to build 3D models, predict ligand binding sites and analyse the effect of amino acid variants on the submitted protein sequence (Lawrence *et al.,* 2015). The intensive mode attempts to create a complete full-length model of a sequence through a combination of multiple template modelling and simplified *ab initio* folding simulation (Lawrence *et al.,* 2015). Three templates were selected to model the final protein model (Figure 5) based on heuristics that maximise confidence, percentage identity and alignment coverage. Two of the templates; c5e4v (PDB Title: crystal structure of measles n0-p complex) and c4uftB (PDB Title: structure of the helical measles virus nucleocapsid) were modelled with 100% confidence. These templates had 81% and 85% identity respectively with the final model.

**Figure 5.**
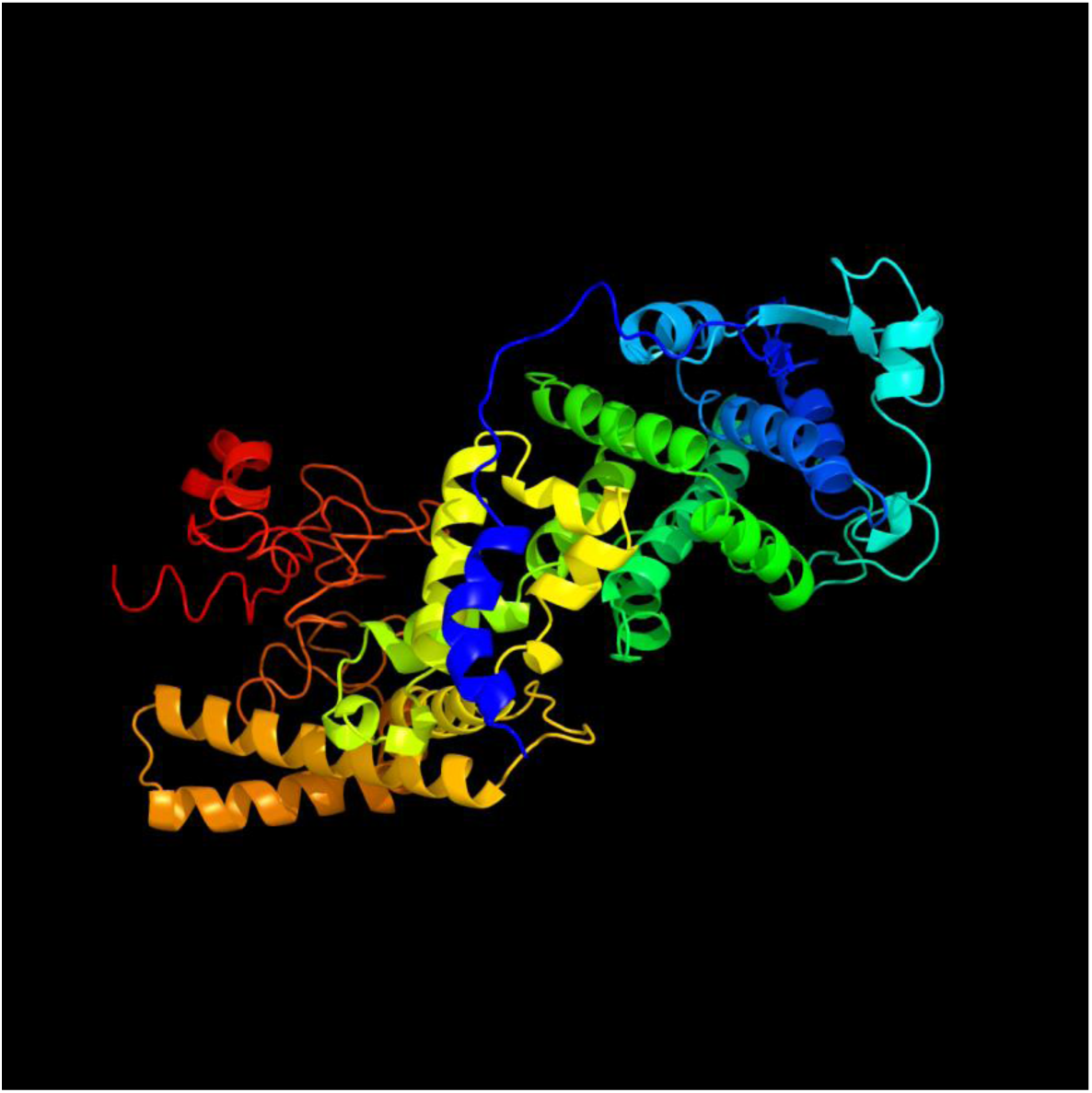
Cartoon structure of the final 3D model of PPRV N predicted by Phyre2.

The third template, c1t6oB (PDB Title: nucleocapsid-binding domain of the measles virus p protein2 (amino acids 457-507) in complex with amino acids 486-5053 of the measles virus n protein) was modelled with 78% confidence, and 80% identity. The confidence of the predicted final model is quite high, with 414 residues (79%) modelled at >90% accuracy. However, 94 residues were modelled by *ab initio.* Ab initio modelling, even though highly unreliable, enables modelling of entire model. Models with a confidence value >90% are considered good quality models. The predicted model is taken to adopt the overall fold shown and that the core of the protein is modelled at an accuracy of 2–4 Å root mean square deviation (RMSD) from the native, true structure (Lawrence *et al.,* 2015).

In addition to the 3D model, ligand binding site of the predicted structure was determined using the 3DLigandSite web server (Wass *et al.,* 2010) and viewed in Jmol. The predicted binding site occurs between amino acid residues ^376^VAL to ^440^ALA of the structure and is shown in blue colour while the rest of the protein is shown as a grey cartoon (Figure 6A). A close up of the ligand binding site (Figure 6B) indicates the residues predicted to form part of the binding site comprising ^376^VAL, ^377^SER, ^380^ILE, ^381^ALA, ^401^THR, ^405^ARG, ^408^ARG, ^412^PRO, ^428^GLU, ^429^SER, ^430^PRO, ^434^THR, ^435^ARG, ^436^GLU, ^437^GLU, ^438^VAL, ^439^LYS and ^440^ALA. The ligands that form the cluster used for the prediction are displayed with non metal ions shown as wireframes.

**Figure 6.**
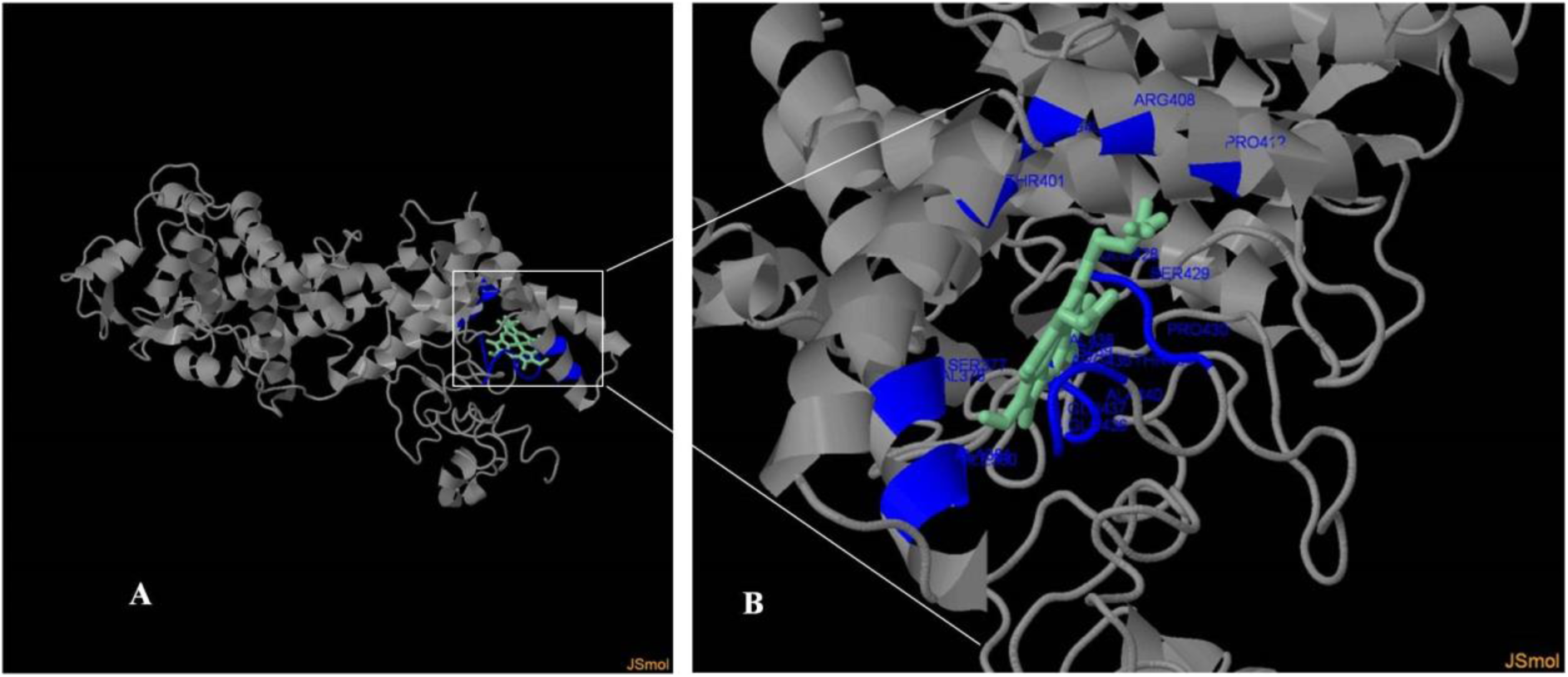
Ligand binding site for predicted 3D PPRV N model.

### Protein structure quality and validation

Integrity of the 3D model was tested with the SAVES server (http://services.mbi.ucla.edu/SAVES/) to obtain ERRAT, VERIFY 3D and PROVE scores. The ERRAT program differentiates between correctly and incorrectly determined regions of protein structures based on characteristic atomic interactions (Colovos and Yeates, 1993). Analysis of frequency distribution and positioning of amino acids in the PPRV N model resulted in a low ERRAT score of 60.665% (Figure 7A). VERIFY 3D Determines the compatibility of an atomic model (3D) with its own amino acid sequence (1D) by assigning a structural class based on its location and environment (Bowie *et al.,* 1991). VERIFY 3D results (Figure 7B) showed that 73.71% of the residues had an averaged 3D-1D score >= 0.2. This score is more than the low 65% mark, but less than the accepted 80% of residues having a 3D-1D score above 0.2. The Z-score of a protein is the difference in energy between the native fold and the average of an ensemble of misfolds in the units of the standard deviation of the ensemble (Zhang and Skolnick, 1998). Z-score for PPRV N (Figure 7C) was predicted with the PROVE (PROtein Volume Evaluation) program (Pointius *et al.,* 1996). The Z-score locates the modelled structure in relation to highly resolved and well-refined protein structures submitted to the PDB (Pointius *et al.,* 1996). The Z-score for PPRV N was found to be −0.095 indicating acceptable model quality. Ramachandran plots (Figure 8) of PPRV N model generated using the RAMPAGE web tool (Lovell *et al.,* 2003)http://mordred.bioc.cam.ac.uk/˜rapper/rampage.php) revealed that the phi-psi torsion angles for 87% of residues are in the favoured region, while 8.4% of the residues are in the allowed region. The number of residues in outlier region was found to be 4.6%. The generally low model quality scores may be as a result of low sequence coverage from templates used for modelling and from lack of PPRV crystal structures solved at near atomic resolution in the PDB.

**Figure 7.**
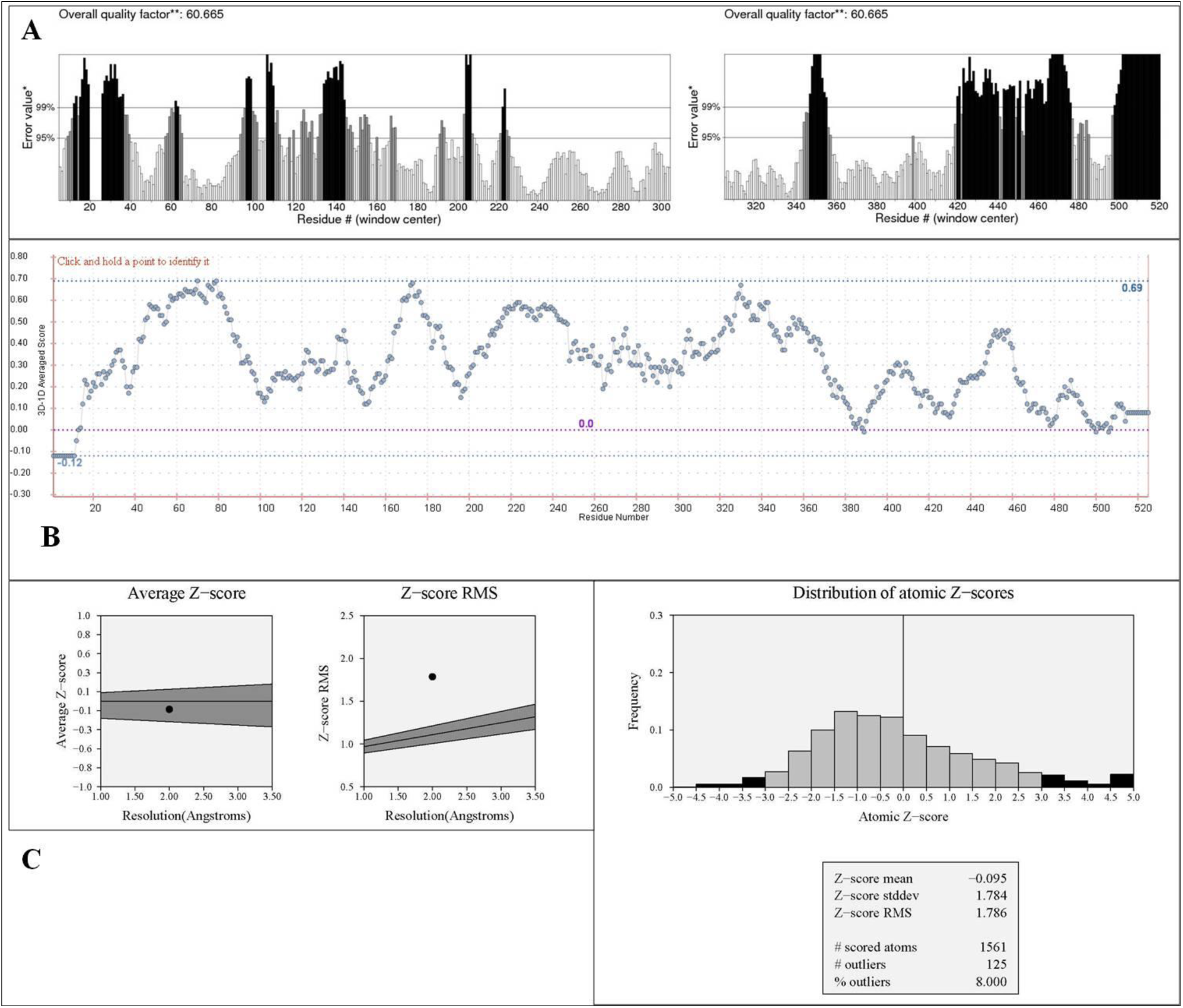
Model quality analysis and verification through. A) ERRAT B) VERIFY 3D and C) PROVE for the 3D PPRV N model

**Figure 8.**
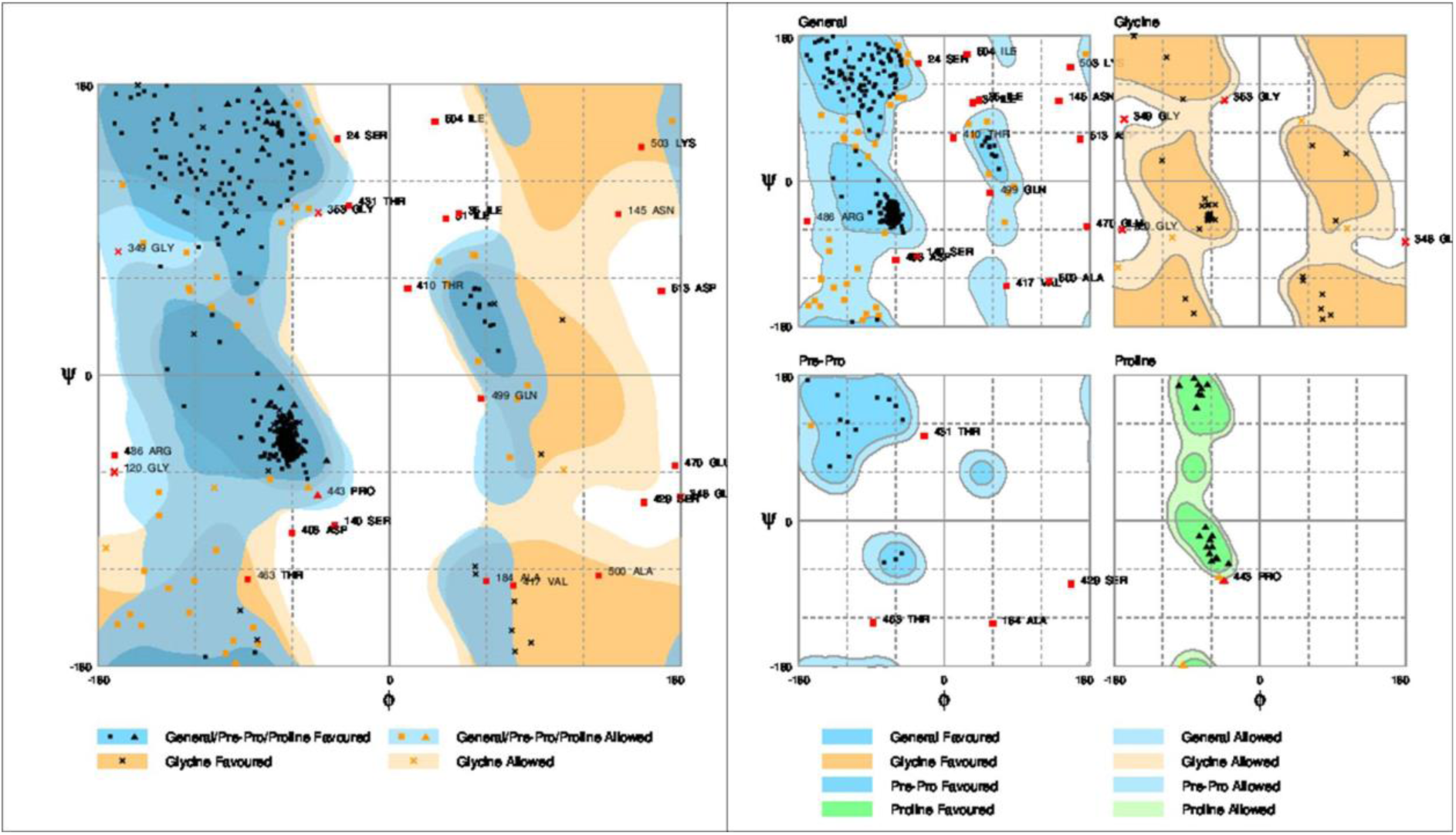
Ramachandran plots of the predicted PPRV N model.

## Conclusion

PPRV is the causal agent of peste des petits ruminants; a highly contagious viral disease with high mortality in small ruminants. The nucleoprotein of PPRV was subjected to epitope prediction using different *in silico* prediction and 3D modelling tools. Predicted epitopes compared favourably to previously described PPRV N epitopes in the literature. Accurate prediction of epitopes is an important part of designing N-protein specific diagnostic immunoassays for PPRV. The next step after *in silico* prediction would be experimental validation of the predicted epitopes and selection of promising candidates for consideration as antigen-based diagnostic tools. Such diagnostic tools would play a role in the global fight and possible eradication of PPR.

## References

Bailey D, Banyard A, Dash P, Ozkul A and Barrett T (2005). Full genome sequence of peste des petits ruminants virus, a member of the *Morbillivirus* genus. Virus Research, 110 (1); 119–124.

Berhe G, Minet C, Le Goff C, Barrett T, Ngangnou A, Grillet C, Libeau G, Fleming M and Diallo A (2003). Development of a dual recombinant vaccine to protect small ruminants against peste-des-petits-ruminants virus and capripoxvirus infections. Journal of Virology, 77(2); 1571–1577.

Bodjo SC, Kwiatek O, Diallo A, Albina E and Libeau G (2007). Mapping and structural analysis of B-cell epitopes on the morbillivirus nucleoprotein amino terminus. Journal of General Virology, 88(4); 1231–1242.

Bowie J, Luthy R and Eisenberg D (1991). A method to identify protein sequences that fold into a known three-dimensional structure. Science, 253(5016); 164–170. (DOI: 10.1126/science.1853201).

Chen J, Liu H, Yang J and Chou K (2007). Prediction of linear B-cell epitopes using amino acid pair antigenicity scale. Amino Acids, 33; 423–428.

Choi KS, Nah JJ, Ko YJ, Kang SY, Yoon KJ and Joo YS (2004). Characterization of immunodominant linear B-cell epitopes on the carboxy terminus of the rinderpest virus nucleocapsid protein. Clinical and Diagnostic Laboratory Immunology, 11(4); 658–664.

Choi SK, Nah JJ, Ko YJ, Kang SY, Yoon KJ and Jo NI (2005). Antigenic and immunogenic investigation of B-cell epitopes in the nucleocapsid protein of Peste des petits ruminants virus. Clinical and Diagnostic Laboratory Immunology, 12(1); 114–121.

Chou PY and Fasman GD (1979). Prediction of beta-turns. Biophysical Journal, 26(3); 367–383.

Colovos C and Yeates TO (1993). Verification of protein structures: patterns of nonbonded atomic interactions. Protein Science, 2(9);1511–1519.

Couacy-Hymann E, Roger F, Hurard C, Guillou JP, Libeau G and Diallo A (2002). Rapid and sensitive detection of peste des petits ruminants virus by a polymerase chain reaction assay. Journal of Virological Methods; 100 (1–2); 17–25

Dechamma HJ, Dighe V, Kumar CA, Singh RP, Jagadish M and Kumar S. (2006). Identification of T-helper and linear B epitope in the hypervariable region of nucleocapsid protein of PPRV and its use in the development of specific antibodies to detect viral antigen. Veterinary Microbiology, 118(3), 201–211.

Dhanda, SK, Usmani, SS, Agrawal P, Nagpal G, Gautam A and Raghava GP (2016). Novel *in silico* tools for designing peptide-based subunit vaccines and immunotherapeutics. Briefings in Bioinformatics. DOI: 10.1093/bib/bbw025.

Diallo A, Barrett T, Barbron M, Meyer G and Lefèvre PC (1994). Cloning of the nucleocapsid protein gene of peste-des-petits-ruminants virus: relationship to other morbilliviruses. Journal of General Virology, 75; 233–237.

EL-Manzalawy, Y, Dobbs, D and Honavar V (2008). Predicting linear B-cell epitopes using string kernels. Journal of molecular recognition, 21(4); 243–255.

Emini EA, Hughes JV, Perlow D and Boger J (1985). Induction of hepatitis A virus-neutralizing antibody by a virus-specific synthetic peptide. Journal of Virology, 55(3); 836–839.

Hoof I, Peters B, Sidney J, Pedersen LE, Sette A, Lund O, Buus S and Nielsen M (2009). NetMHCpan, a method for MHC class I binding prediction beyond humans. Immunogenetics, 61(1); 1–13.

Hopp TP and Woods KR (1981). Prediction of protein antigenic determinants from amino acid sequences. Proceedings of the National Academy of Sciences, 78(6); 3824–3828.

Ismail TM, Yamanaka MK, Saliki JT, El-Kholy A, Mebus C and Yilma T (1995). Cloning and expression of the nucleoprotein of Peste des petits ruminants virus in Baculovirus for use in serological diagnosis. Virology 208, 776–778

Karplus PA and Schulz, GE (1985). Prediction of chain flexibility in proteins. Naturwissenschaften, 72(4); 212–213.

Kelley LA, Mezulis S, Yates CM, Wass MN and Sternberg MJ (2015). The Phyre2 web portal for protein modeling, prediction and analysis. Nature Protocols, 10(6); 845–858.

Keşmir C, Nussbaum, AK, Schild H, Detours V and Brunak S (2002). Prediction of proteasome cleavage motifs by neural networks. Protein Engineering, 15(4); 287–296.

Kolaskar, AS and Tongaonkar PC (1990). A semi-empirical method for prediction of antigenic determinants on protein antigens. FEBS Letters, 276(1-2); 172–174.

Kringelum JV, Lundegaard C, Lund O, Nielsen M (2012). Reliable B cell epitope predictions: impacts of method development and improved benchmarking. PLOS Computational Biology, 8(12); e1002829. DOI:10.1371/journal.pcbi.1002829.

Kyte J and Doolittle RF (1982). A simple method for displaying the hydropathic character of a protein. Journal of Molecular Biology, 157(1); 105–132.

Larsen MV, Lundegaard C, Lamberth K, Buus S, Brunak S, Lund O and Nielsen M (2005). An integrative approach to CTL epitope prediction: a combined algorithm integrating MHC class I binding, TAP transport efficiency, and proteasomal cleavage predictions. European Journal of Immunology, 35(8); 2295–2303.

Larsen JEP, Lund O and Nielsen M (2006). Improved method for predicting linear B-cell epitopes. Immunome Research, 2(2). http://www.immunome-research.com/content/2/1/2

Larsen MV, Lundegaard C, Lamberth K, Buus S, Lund O and Nielsen M (2007). Large-scale validation of methods for cytotoxic T-lymphocyte epitope prediction. BMC Bioinformatics, 8(424). http://doi.org/10.1186/1471-2105-8-424

Lovell SC, Davis IW, Arendall WB, de Bakker PIW, Word JM, Prisant MG, Richardson JS and Richardson DC (2003). Structure validation by Cα geometry: ϕ,ψ and Cβ deviation. Proteins, 50; 437–450. DOI: 10.1002/prot.10286.

Lundegaard C, Lamberth K, Harndahl M, Buus S, Lund, O and Nielsen M (2008). NetMHC-3.0: accurate web accessible predictions of human, mouse and monkey MHC class I affinities for peptides of length 8–11. Nucleic Acids Research, 36(suppl 2); W509–W512.

McWilliam H, Li W, Uludag M, Squizzato S, Park YM, Buso N, Cowley AP and Lopez R (2013). Analysis tool web services from the EMBL-EBI. Nucleic Acids Research, 41(W1); W597–W600. DOI:10.1093/nar/gkt376.

Michalik M, Djahanshiri B, Leo JC and Linke D (2016). Reverse vaccinology: the pathway from genomes and epitope predictions to tailored recombinant vaccines. In, Thomas S (Ed). Vaccine Design: Methods and Protocols: Volume 1: Vaccines for Human Diseases. Springer Science, New York. pp. 87–106.

Mitra-Kaushik S, Nayak R and Shaila MS (2001). Identification of a Cytotoxic T-Cell Epitope on the recombinant nucleocapsid proteins of Rinderpest and Peste des petits ruminants viruses presented as assembled nucleocapsids. Virology, 279; 210–220.

Nielsen M and Andreatta M (2016). NetMHCpan-3.0; improved prediction of binding to MHC class I molecules integrating information from multiple receptor and peptide length datasets. Genome Medicine, 8(33). http://doi.org/10.1186/s13073-016-0288-x

Nielsen M and Lund O (2009). NN-align. An artificial neural network-based alignment algorithm for MHC class II peptide binding prediction. BMC Bioinformatics, 10(1). DOI: 10.1186/1471-2105-10-296

Nielsen M, Lundegaard C, Blicher T, Peters B, Sette A, Justesen S, Buus S and Lund O (2008). Quantitative predictions of peptide binding to any HLA-DR molecule of known sequence: NetMHCIIpan. PLOS Computational Biology, 4(7); p. e1000107.

Nielsen M, Lundegaard C and Lund O (2007). Prediction of MHC class II binding affinity using SMM-align, a novel stabilization matrix alignment method. BMC Bioinformatics, 8(1). DOI: 10.1186/1471-2105-8-238.

Nielsen M, Lundegaard C, Lund O and Keşmir C (2005). The role of the proteasome in generating cytotoxic T-cell epitopes: insights obtained from improved predictions of proteasomal cleavage. Immunogenetics, 57(1-2); 33–41.

Parker JMR, Guo D and Hodges R.S (1986). New hydrophilicity scale derived from high-performance liquid chromatography peptide retention data: correlation of predicted surface residues with antigenicity and X-ray-derived accessible sites. Biochemistry, 25(19); 5425–5432.

Pontius J, Richelle J and Wodak SJ (1996). Deviations from standard atomic volumes as a quality measure for protein crystal structures. Journal of Molecular Biology, 264(1); 121–136.

Ponomarenko JV and Van Regenmortel MH (2009). B cell epitope prediction. In Gu J and Bourne PE (Eds). Structural Bioinformatics, Second Edition. Hoboken, NJ; Wiley-Blackwell. pp. 849–879.

Ponomarenko J, Bui HH, Li W, Fusseder N, Bourne PE, Sette A and Peters B (2008). ElliPro: a new structure-based tool for the prediction of antibody epitopes. BMC Bioinformatics, 9(1). DOI: 10.1186/1471-2105-9-514.

Pope RA, Parida S, Bailey D, Brownlie J, Barrett T and Banyard AC (2013). Early events following experimental infection with Peste-Des-Petits ruminants virus suggest immune cell targeting. PLOS One, 8(2), e55830.

Saxová P, Buus S, Brunak S and Keşmir C (2003). Predicting proteasomal cleavage sites: a comparison of available methods. International Immunology, 15(7); 781–787.

Sette A, Livingston B, McKinney D, Appella E, Fikes J, Sidney J, Newman M and Chesnut R (2001). The development of multi-epitope vaccines: epitope identification, vaccine design and clinical evaluation. Biologicals, 29(3); 271–276.

Shaila MS, Shamaki D, Forsyth MA, Diallo A, Goatley L, Kitching RP and Barrett T (1996). Geographic distribution and epidemiology of peste des petits ruminants viruses. Virus Research; 43 (2), 149–153.

Stranzl T, Larsen MV, Lundegaard C and Nielsen M (2010). NetCTLpan: pan-specific MHC class I pathway epitope predictions. Immunogenetics, 62(6); 357–368.

Sturniolo T, Bono E, Ding J, Raddrizzani L, Tuereci O, Sahin U, Braxenthaler M, Gallazzi F, Protti MP, Sinigaglia F and Hammer J (1999). Generation of tissue-specific and promiscuous HLA ligand databases using DNA microarrays and virtual HLA class II matrices. Nature Biotechnology, 17(6); 555–561.

Sun P, Ju H, Zhang B, Gu Y, Liu B, Huan Y, Zhang H and Li Y (2015). Conformational B-cell epitope prediction method based on antigen preprocessing and mimotopes analysis. BioMed Research International, 2015. http://dx.doi.org/10.1155/2015/257030.

Tong JC, Tan TW and Ranganathan S (2007). Methods and protocols for prediction of immunogenic epitopes. Briefings in Bioinformatics, 8(2); 96–108.

Wang P, Sidney J, Dow C, Mothè B, Sette A and Peters B (2008). A systematic assessment of MHC class II peptide binding predictions and evaluation of a consensus approach. PLOS Computational Biology, 4(4); e1000048. DOI: 10.1317/journal.pcbi.1000048.

Wang P, Sidney J, Kim Y, Sette A, Lund O, Nielsen M and Peters B (2010). Peptide binding predictions for HLA DR, DP and DQ molecules. BMC Bioinformatics, 11(1). DOI: 10.1186/1471-2105-11-568.

Wass MN, Kelley LA and Sternberg MJE (2010). 3DLigandSite: predicting ligand-binding sites using similar structures. Nucleic Acids Research, 38(Web Server issue), W469–W473. http://doi.org/10.1093/nar/gkq406

Yadav V, Balamurugan V, Bhanuprakash V, Sen A, Bhanot V, Venkatesan G, Riyesh T, Singh RK (2009). Expression of Peste des petits ruminants virus nucleocapsid protein in prokaryotic system and its potential use as a diagnostic antigen or immunogen. Journal of Virological Methods 162, 56–63.

Yu R, Fan X, Xu, W, Li W, Dong S, Zhu Y, He Y, Tang H, Du R and Li Z (2015). Fine mapping and conservation analysis of linear B-cell epitopes of peste des petits ruminants virus nucleoprotein. Veterinary Microbiology, 175(1); 132–138.

Zepp F (2016). Principles of vaccination. In Thomas S (Ed). Vaccine Design: Methods and Protocols: Volume 1: Vaccines for Human Diseases. Springer Science, New York. pp. 57–84.

Zhang L and Skolnick J (1998). What should the Z-score of native protein structures be? Protein Science, 7; 1201–1207.

